# Optogenetic stimulation of lateral hypothalamic orexin/dynorphin inputs in the ventral tegmental area potentiates mesolimbic dopamine neurotransmission and promotes reward-seeking behaviours

**DOI:** 10.1101/2020.09.10.291963

**Authors:** Catherine S. Thomas, Aida Mohammadkhani, Madiha Rana, Min Qiao, Corey Baimel, Stephanie L. Borgland

## Abstract

Reward and reinforcement processes are critical for survival and propagation of genes. While numerous brain systems underlie these processes, a cardinal role is ascribed to mesolimbic dopamine. However, ventral tegmental area (VTA) dopamine neurons receive complex innervation and various neuromodulatory factors, including input from lateral hypothalamic (LH) orexin/hypocretin neurons which also express and co-release the neuropeptide, dynorphin. Dynorphin in the VTA induces aversive conditioning through the Kappa opioid receptor (KOR) and decreases dopamine when administered intra-VTA. Exogenous application of orexin or orexin 1 receptor (oxR1) antagonists in the VTA bidirectionally modulates dopamine-driven motivation and reward-seeking behaviours, including the attribution of motivational value to primary rewards and associated conditioned stimuli. However, the effect of endogenous stimulation of LH orexin/dynorphin-containing projections to the VTA and the potential contribution of co-released dynorphin on mesolimbic dopamine and reward related processes remains uncharacterised. We combined optogenetic, electrochemical, and behavioural approaches to examine this. We found that optical stimulation of LH orexin/dynorphin inputs in the VTA potentiates mesolimbic dopamine neurotransmission in the nucleus accumbens (NAc) core, produces real time and conditioned place preference, and increases the food cue-directed orientation in a Pavlovian conditioning procedure. LH orexin/dynorphin potentiation of NAc dopamine release and real time place preference was blocked by an oxR1, but not KOR antagonist. Thus, rewarding effects associated with optical stimulation of LH orexin/dynorphin inputs in the VTA are predominantly driven by orexin rather than dynorphin.

## INTRODUCTION

Ventral tegmental area (VTA) dopamine neurons are involved in motivated survival behaviours and their activity can influence positive and negative reinforcement, incentive salience, and aversion[1–4]. These varied functions are guided by subpopulations of VTA dopamine neurons [5], efferent projections to different regions [6–8], different neuromodulatory afferents [5,9–11], or a combination of these factors. Mesolimbic dopamine projections from the VTA to the nucleus accumbens (NAc) energize behaviour toward motivationally relevant cues[12,13]. However, factors that modulate dopamine in relation to motivated behaviour are complex and not fully understood.

One factor that modulates mesolimbic dopamine neuronal activity, plasticity and reward-seeking is the lateral hypothalamic (LH) peptide, orexin (or hypocretin [14]). Orexin A and B are synthesized in the LH area and project to cortical and subcortical brain structures [15–17]. The VTA receives en passant LH orexin-containing fibres with close appositions to VTA dopamine neurons [15,18,19]. Although these fibres are present in the VTA, less than 5% show identifiable synaptic specializations [18]. However, most axons in the VTA stain heavily for dense core vesicles[18], suggesting that, similar to other neuropeptides, orexin release in the VTA is largely extrasynaptic. LH orexin-containing neurons are highly co-localized with dynorphin [20], the endogenous ligand of kappa opioid receptors (KOR). Co-expression is observed in dense core vesicles, suggesting that these peptides may be co-released [21,22]. Indeed, co-release of orexin and dynorphin within the LH have opposing actions on firing of local neurons [22]. Similarly, exogenous orexin application to VTA slices increases firing of dopamine neurons projecting to the NAc shell, but not the basolateral amygdala, whereas bath application of dynorphin has the opposite effect, suggesting a distinct role for orexin in promoting VTA-NAc dopamine activity [10]. While orexin in the VTA augments reward-seeking and motivated behaviours through activation of orexin receptor 1 (oxR1) [23–27], dynorphin in the VTA inhibits reward-seeking and produces conditioned place aversion [28,29] and reduced dopamine [30]. OxR1 antagonism in the VTA decreases cocaine self-administration and LH-self-stimulation, both which are reversed by blocking KORs. Thus, it has been proposed that orexin signaling can occlude the reward threshold-elevating effects of co-released dynorphin and thereby facilitate reward-seeking [21]. While LH orexin inputs provide the only source of orexin to the VTA, several brain nuclei have dynorphin-containing neurons [31,32], with significant dynorphin-containing projections from the NAc to the VTA [33]. Because application of intra-VTA norBNI inhibits actions of dynorphin from all sources, it is unknown what the direct contribution of dynorphin from the LH input is to reward-seeking behaviour. Furthermore, it is unknown if endogenous LH orexin/dynorphin action in the VTA can influence mesolimbic dopamine neurotransmission and motivational processes that guide reward-seeking. In this study, we systematically tested whether optical stimulation of LH orexin/dynorphin afferents in the VTA promotes NAc dopamine release, reward-seeking behaviours, and/or increases the incentive value of food.

## BRIEF METHODS

### Detailed methods are located in the supplemental material

#### Subjects

Adult male and female orexin-EGFP-2A-cre (orexin-cre) mice (post-natal day 60-90; originally from the Yamanaka lab at the Nagoya University [34]) were bred locally at the University of Calgary. All mice received bilateral infusions of either channelrhodopsin (‘ChR2’) (AAV2/8-EF1a-DIO-hChR2(H134R)-mCherry) or control (‘mCherry’) (AAV2/8-hSyn-DIO-mCherry) virus. All experimental procedures adhered to ethical guidelines established by the Canadian Council for Animal Care and animal use protocols approved by the University of Calgary Animal Care Committee.

#### Real-Time Place Preference (RTPP)

Orexin-cre ChR2 (n=8) and Orexin-cre mCherry (n=10) mice implanted with bilateral optical fibres 6 weeks after viral infusions underwent 3 stages of real-time place preference paradigm. Briefly, to assess baseline preference, mice explored a 2-compartment arena while receiving no stimulation. Next, mice underwent 3 days of stimulation sessions where one compartment (counterbalanced across mice) was paired with optical stimulation of intra-VTA LH orexin/dynorphin (473 nm, 20 Hz, 5 mW, 5 ms pulses, 1 s duration). Lastly, on the test day, mice explored the compartments with no stimulation.

For experiments involving the oxR1 antagonist, SB-334867, or the KOR antagonist, nor BNI, orexin-cre mice were infused with ChR2 (n = 10 for SB-334867 and n = 12 for norBNI) in LH and implanted with optical fibres targeted at the VTA. Mice underwent RTPP as above. Prior to each conditioning session (day 2-4), all mice received either SB-334867 (n = 5, 15 mg/kg dissolved in 10% hydroxypropyl-beta-cyclodextrin and 2% dimethyl sulfoxide (DMSO) in sterile water w/v) or vehicle (n = 5), or norBNI (n = 5, 15 mg/kg in saline) or vehicle (n = 7) 15 min prior to being placed into the arena.

#### Pavlovian Conditioning

Orexin-Cre ChR2 (n=8) or mCherry (n=10) mice implanted with bilateral optical fibres targeted at the VTA were used for both RTPP and Pavlovian conditioning in a counter-balanced order. Mice underwent baseline testing to examine individual differences in value of novel neutral stimulus and sucrose pellets prior to conditioning. Next all mice underwent 7 consecutive days of Pavlovian conditioning, where delivery of a sucrose pellet into the behavioural chamber was paired with LH orexin/dynorphin intra-VTA optical stimulation, and 8 s cue presentation (light + tone; see detailed description in supplemental materials). After conditioning, all mice underwent conditioned reinforcement to examine any change in individual responding for the food cue. Lastly, all mice were given sucrose pellets in the home cage to re-assess the value of the primary reward after conditioning, compared to individuals baseline levels.

All Pavlovian conditioning sessions were video recorded. The videos were then scored for approach and orientation (including latency to orient and latency to approach) to the cue by experimenters who were blinded to experimental conditions. Videos were each scored twice to provide inter-rater reliability.

#### *In vivo* Fast-Scan Cyclic Voltammetry (FSCV)

Mice (ChR2 n= 7; mCherry n=7) were anaesthetized with 25% urethane (1.0-1.5 g/kg in saline). Small craniotomies were made above the NAc (AP +1.0; ML +1.0) and the VTA (AP -3.5; ML +0.5) and contralateral cortex (AP +1.8; ML -2.0). A chlorinated silver wire reference electrode was implanted in the contralateral cortex (DV -0.8) and cemented in place. Recording (targeted at right NAc core) and stimulating (combined with optical fibre; targeted at VTA) were slowly lowered to desired locations. FSCV data were collected in 120 s files, with stimulation onset occurring 5 s into the recording using TarHeel CV and High-Definition Cyclic Voltammetry (HDCV) Suite. 10 recordings of electrical stimulation (60 Hz, 60 pulses; 120 μA), 10 recordings of optical stimulation (laser as above; 473 nm, 20 Hz, 20mW, 5-ms pulses, 1s duration), and 10 recordings of combined electrical and optical stimulation were made in a counter-balanced order across subjects.

For SB-334867 (ChR2 n = 7; mCherry n = 7) or norBNI (ChR2 n = 6; mCherry n = 5) injections, we conducted anaesthetised FSCV data acquisition as above, followed by SB-334867 (15 mg/kg) or norBNI (15 mg/kg). After 15 min, we recorded evoked responses to electrical stimulation and to electrical + optical stimulation and compared the area under the curve and peak dopamine concentration to recordings made prior to administration of the antagonist.

#### Statistics

All statistical analyses were completed using GraphPad Prism 8 or 9. All values are expressed as mean ± SEM. The alpha risk for the rejection of the null hypothesis was set to 0.05. All data met criteria for normality unless otherwise specified. All post hoc tests were conducted with Sidak’s correction for multiple comparisons, unless otherwise stated. *P<0.05, **P<0.01, P<0.001***, P<0.0001***.

## RESULTS

We first determined the window of efficacy for optical control of ChR2-expressing LH orexin/dynorphin neuronal activity ex vivo. Using whole-cell patch recordings, we confirmed the selectivity of the floxed-AAV-cre strategy to genetically target the expression of ChR2 to LH orexin/dynorphin neurons and the reliability of photocurrents in ChR2-expressing LH orexin/dynorphin neurons over a wide range of frequencies (1-100 Hz). LH orexin/dynorphin neurons can be reliably activated by photostimulation (Supplemental Figure 1A-D). We next identified the frequency of optical stimulation of LH orexin/dynorphin inputs that would influence firing activity of VTA dopamine neurons (Supplemental Figure 1E). A 20 Hz stimulation increased firing of dopamine neurons and we used this frequency for in vivo experiments (Supplemental Figure 1F).

### Endogenous LH orexin/dynorphin release in the VTA promotes place preference

To examine the hypothesis that optogenetic stimulation of LH orexin/dynorphin inputs in the VTA would promote place preference (Figure 1A-D), we used a real-time place preference (RTPP) procedure, whereby time spent in one of the compartments is paired with intra-VTA optogenetic stimulation of LH orexin/dynorphin inputs. In the absence of stimulation, there was no significant baseline preference for either compartment displayed by mCherry control or ChR2 expressing mice (Figure 1E; mCherry: *t*(7) = 1.6, *p* = 0.2; ChR2: *t*(7) = 0.8, *p* = 0.5).Furthermore, mCherry mice did not develop a significant preference for either compartment of the apparatus over the 3 RTPP stimulation sessions (Figure 1E). However, the ChR2 mice demonstrated a significant preference for the compartment paired with intra-VTA LH orexin/dynorphin stimulation over the 3 stimulation days (Figure 1E,F; 3 × 2 RM ANOVA: Day, F(2,32) = 6.8; *p* = 0.004; Virus, F(1, 16) = 0.15, *p* = 0.5; Compartment, F(1,16) = 17.2, *p* = 0.0008; Day x Virus: F(2,32) = 0.7, *p* = 0.5; Day x Compartment: F(2,32) = 5.7, *p* = 0.008; Virus x Compartment, F(1,16) = 19.8, *p* = 0.0004; Day x Compartment x Virus, F(2, 32) = 2.5, *p* = 0.09). ChR2 mice spent significantly more time in the stimulation compartment compared to the non-stimulation compartment on days 2 and 3 (Sidak’s post hoc: day 2: *t*(7) = 3.4; *p* = 0.01; day 3: *t*(7) = 4.2, *p* = 0.004).

**Figure 1.**
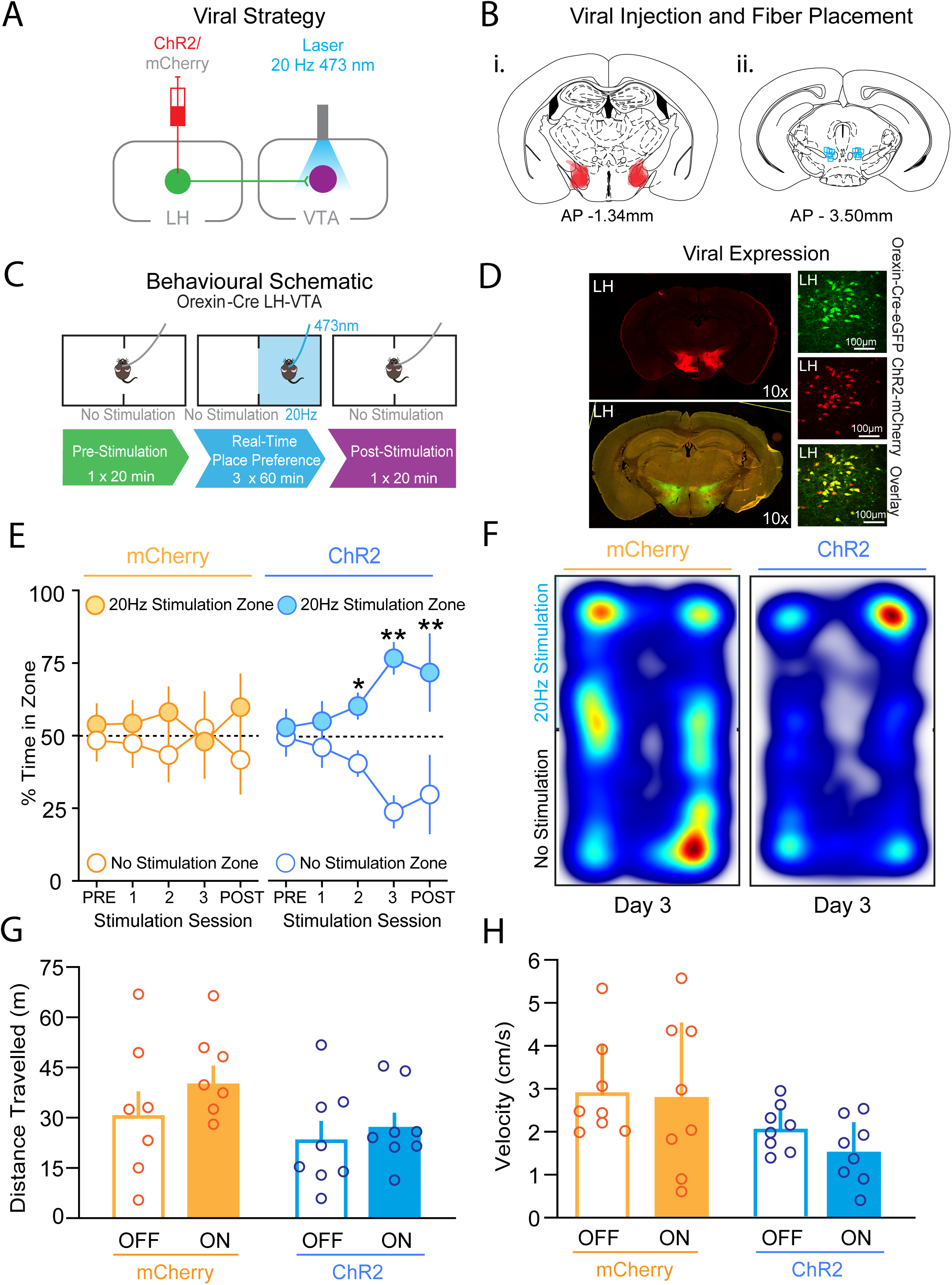
Optogenetic stimulation of LH orexin/dynorphin inputs in the VTA promotes place preference. A) Viral strategy schematic. Orexin-cre mice were infused with either channelrhodopsin (AAV2/8-EF1a-DIO-hChR2(H134R)-mCherry; ‘ChR2’) or control virus (AAV2/8-hSyn-DIO-mCherry; ‘mCherry’). (LH: lateral hypothalamus; VTA: ventral tegmental area) B) Coronal brain sections (modified from [55]) representing i) estimated viral transfection areas in the LH (−1.34 mm in anterior-posterior plane relative to Bregma) and ii) estimated location of optical fibre implantation site in the VTA (−3.50 mm in anterior-posterior plane relative to Bregma). C) Schematic of behavioural apparatus used for all real-time place preference (RTPP) experiments. Blue square in middle panel denotes the chamber associated with 473 nm laser stimulation of LH orexin/dynorphin. Compartments were counterbalanced across mice. D) Immunohistochemical demonstration of fluorophore expression in LH cell bodies genetically encoded fluorophore (green; orexin-cre-eGFP) and viral transfection fluorophore (red; ChR2 mCherry); ChR2 virus infected cells were colocalized with orexin. E) Time course of RTPP prior to and during optogenetic stimulation of LH orexin/dynorphin inputs in the VTA of mCherry (orange) or ChR2 (blue) mice in the compartment paired with stimulation (filled circles) and the no stimulation compartment (open circles). Over 3 stimulation sessions of RTPP, ChR2 mice developed a significant preference for the compartment paired with LH orexin/dynorphin stimulation whereas mCherry mice did not. In the post-stimulation test session, ChR2 mice preferred the compartment previously paired with stimulation, whereas mCherry mice did not. F) Representative heat maps of percent time spent in both compartments of the RTPP arena for mCherry and ChR2 mice on stimulation session 3. G) Optical stimulation of LH orexin/dynorphin inputs to the VTA did not alter the distance travelled by mCherry and ChR2 mice in either the stimulation ON (filled bars) or stimulation OFF (open bars) compartments of the RTPP arena during stimulation sessions. H) Optical stimulation of LH orexin/dynorphin inputs to the VTA did not alter velocity of locomotor activity mCherry and ChR2 mice in either the stimulation ON (filled bars) or stimulation OFF (open bars) compartments during stimulation sessions.

To determine if intra-VTA photostimulation of LH orexin/dynorphin can influence associative learning mechanisms, we next examined whether mCherry or ChR2 mice demonstrated a preference for either compartment in the absence of optical stimulation in a subsequent session. While the mCherry mice did not show a significant preference for either compartment, the ChR2 mice spent significantly more time in the compartment previously paired with stimulation (Figure 1E; Sidak’s post hoc: mCherry: *t*(7) = 0.7, *p* = 0.5; ChR2: *t*(7) = 4.6, *p* = 0.002). Thus, LH orexin/dynorphin in the VTA can induce rewarding effects upon stimulation as well as form associative memories related to it.

To assess if locomotor activity was influenced by stimulation, we compared total distance travelled, which did not differ between the mCherry (8.6 ± 1.5 m) and ChR2 (6.0 + 2. 6 m) groups (*t*(14) = 1.7, *p* = 0.1). Velocity also did not differ between mCherry (0.003 ± 0.0005 m/s) and ChR2 (0.002 ± 0.0005 m/s) mice (*t*(7) =1.6, *p* = 0.2). We then examined whether locomotor activity differed between mCherry and ChR2 mice in each compartment. A 2-way ANOVA showed that distance travelled in the stimulation ON or stimulation OFF compartments did not significantly differ (Figure 1G; 2-way ANOVA: Compartment, F(1,28) = 1.4, *p* = 0.2; Virus, F(1, 28) = 3.4, *p* = 0.08; Virus x Compartment, F(1,28) = 0.3, *p* = 0.6). Velocity also did not differ between stimulation ON or stimulation OFF chambers (Figure 1H; 2-way ANOVA: Compartment, F(1,28) = 0.5, *p* = 0.5; Virus, F(1,28) = 6.4, *p* = 0.02; Compartment x Virus: F(1, 28) = 0.2, *p* = 0.7). Thus, intra-VTA optical stimulation of LH orexin/dynorphin produced a preference for the compartment paired with stimulation, without influencing locomotor activity, suggesting that LH orexin/dynorphin in the VTA can promote contingent learning.

As LH orexin/dynorphin-containing neurons co-release the inhibitory neuropeptide dynorphin [20,21] and the fast-acting neurotransmitter glutamate with orexin [35], we first examined the role of oxR1 signalling using the oxR1 antagonist SB-334867, in the RTPP paradigm. As in Figure 1, ChR2 mice did not show a significant preference for either compartment prior to stimulation sessions (Figure 2A; Vehicle: *t*(4)= 0.6, *p* = 0.6; SB-334867: *t*(4)=0.5, *p* = 0.6). Across subsequent conditioning sessions, the vehicle group developed a significant preference for the compartment paired with LH orexin/dynorphin stimulation in the VTA (RM ANOVA: Day, F(4,20)= 4.8, *p* = 0.007; Compartment, F(1,20)= 59.5, *p* < 0.0001; Day x Compartment, F(4,20)= 3.9, *p* = 0.02, Sidak’s post hoc for stimulation on days 1 (*p* = 0.02), 2 (*p* = 0.006), and 3 (*p* < 0.0001) (Figure 2A, B). Whereas, the SB-334867 group, treated prior to each conditioning session, did not develop a significant preference for either compartment (RM ANOVA: Day, F(4,20) = 0.7, *p* = 0.6, Compartment, F(1,20)= 0.004, *p* = 0.9, Day x Compartment, F(4,20) = 0.2, *p* = 0.9). ChR2 mice receiving vehicle injections also demonstrated a preference for the stimulation ON chamber during the post stimulation test session, whereas ChR2 mice receiving SB-334867 did not (Vehicle: *t*(4)= 4.1, *p* =0.002; SB-334867: *t*(4) = 0.2, *p* =0.9) (Figure 2A).

**Figure 2.**
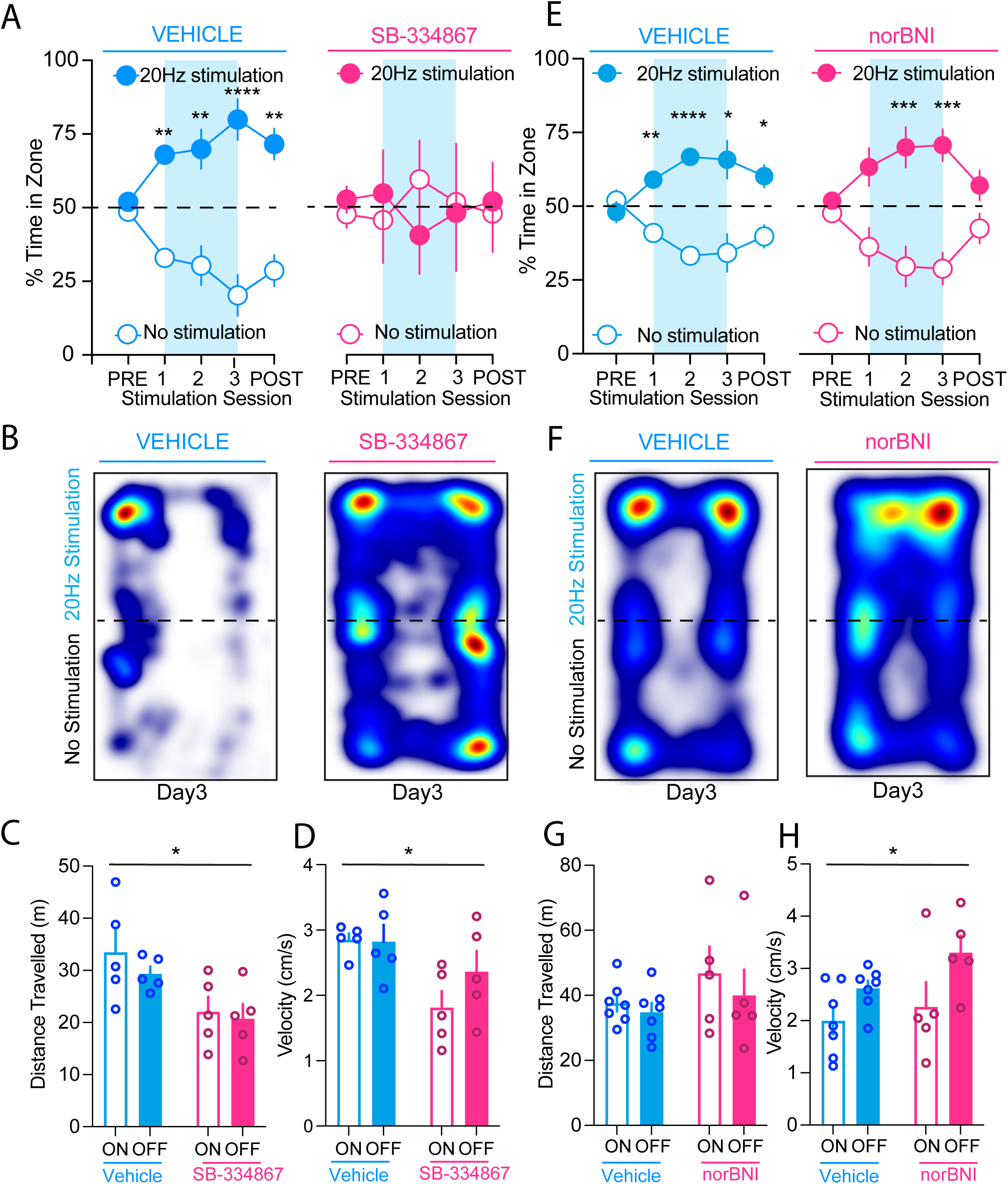
OxR1 mediates preference induced by optogenetic stimulation of LH orexin/dynorphin inputs in the VTA. A) Pre-vehicle (blue) or pre-SB-334867 (pink) mice did not show a significant preference for either compartment prior to stimulation sessions. Over 3 stimulation sessions of RTPP, ChR2 mice receiving vehicle injections developed a preference for the chamber paired with LH orexin/dynorphin stimulation (filled circles) in the VTA, whereas ChR2 mice receiving SB-334867 injections did not develop a preference for the stimulation ON compartment (filled circles) or the stimulation OFF compartment (open circles) of the RTPP arena. ChR2 mice receiving vehicle injections (blue) also demonstrated a preference for the stimulation ON chamber (filled circles) during the post stimulation test session, whereas ChR2 mice receiving SB-334867 (pink) did not. B) Representative heat maps of the percent time spent on day 3 of stimulation sessions in both compartments of the RTPP arena by ChR2 mice receiving either vehicle (left) or SB-334867 (right) injections prior to each stimulation session. C) Distance travelled of ChR2 mice was reduced in mice administered SB-334867 compared to vehicle averaged over the 3 stimulation sessions. However, there was no difference in distance travelled in compartments with intra-VTA LH orexin/dynorphin stimulation ON (filled bars) compared to stimulation OFF (open bars). D) Velocity of ChR2 mice was reduced in mice administered SB-334867 compared to vehicle averaged over the 3 stimulation sessions. However, there was no difference in velocity in compartments with intra-VTA LH orexin/dynorphin stimulation ON (filled bars) compared to stimulation OFF (open bars). E) ChR2 mice receiving vehicle (blue) or norBNI injections (pink) did not show a significant preference for either compartment prior to stimulation sessions. Over 3 stimulation sessions of RTPP, ChR2 mice receiving vehicle or norBNI injections developed a preference for the chamber paired with LH orexin/dynorphin stimulation (filled circles) in the VTA. ChR2 mice receiving vehicle injections also demonstrated a preference for the stimulation ON chamber during the post stimulation test session, whereas ChR2 mice receiving norBNI did not. F) Representative heat maps of the percent time spent on day 3 of stimulation sessions in both compartments of the RTPP arena by ChR2 mice receiving either vehicle (left) or norBNI (right) injections prior to each stimulation session. G) Distance travelled of ChR2 mice was not different in mice administered norBNI compared to vehicle averaged over the 3 stimulation sessions in compartments with intra-VTA LH orexin/dynorphin stimulation ON (filled bars) compared to stimulation OFF (open bars). H) Velocity of ChR2 mice was reduced in mice administered norBNI compared to vehicle averaged over the 3 stimulation sessions. However, there was no difference in velocity in compartments with intra-VTA LH orexin/dynorphin stimulation ON compared to stimulation OFF.

To determine if SB-334867 influenced locomotor activity in either compartment, we measured distance travelled and velocity. There was a significant main effect of SB-334867 on distance travelled, but no interaction between drug and compartment, suggesting that the influence of SB-334867 on locomotor activity did not have differential effects in the stimulation ON or OFF compartment (Figure 2C; 2-way ANOVA: Drug, F(1,8) = 8.6, *p* = 0.02; Compartment, F(1,8) = 1.1, *p* = 0.2; Drug x Compartment, F(1,8) = 0.3, *p* = 0.6). Administration of SB-334867 had a significant effect on velocity, but this was not different between stimulation ON and OFF compartments (Figure 2D; 2-way ANOVA: Drug, F(1,8) =5.7, *p* = 0.04; Compartment, F(1,8) = 3.2, *p* = 0.1, Drug x Compartment, F(1,8)= 3.8, *p* = 0.09).

Lastly, in the post-stimulation test session, mice injected with vehicle spent significantly more time in the paired compartment compared to the unpaired compartment, whereas the mice given SB-334867 showed no significant preference for either compartment (Figure 2A; Vehicle: *t*(6)= 0.6, *p* = 0.6; norBNI: *t*(4)=1.6, *p* = 0.2). Thus, inhibition of oxR1 signalling prevented both the development of preference for LH orexin/dynorphin optical stimulation in the VTA as well as its associated memory.

To examine the contribution of LH dynorphin from optically stimulated LH orexin/dynorphin inputs in the VTA to RTPP, we used the KOR antagonist, norBNI, prior to each conditioning session. ChR2 mice receiving vehicle or norBNI injections did not show a significant preference for either compartment prior to stimulation sessions. Across subsequent conditioning sessions, both the vehicle group and the norBNI group developed a significant preference for the compartment paired with LH orexin/dynorphin stimulation in the VTA (Figure 2E,F) (**Vehicle**: RM ANOVA: Day, F(1.9,22.8)= 0.00, *p* >0.99; Compartment, F(1,12)= 32.4, *p* < 0.0001; Day x Compartment, F(4,48)= 8.3, *p* < 0.0001; **norBNI:** RM ANOVA: Day, F(1.8,14.7) = 0.0004, *p* > 0.99, Compartment: F(1,8)= 32.5, *p* = 0.0005, Day x Compartment, F(4,32) = 5.6, *p* = 0.002). Sidak’s multiple comparisons tests revealed significant differences on days 1 (*p* = 0.004), 2 (*p* < 0.0001), and 3 (*p* = 0.02) in the vehicle group and on days 2 (*p* = 0.02), and 3 (*p* = 0.003) in the norBNI group).

To determine if norBNI influenced locomotor activity in either compartment, we measured distance travelled and velocity. There was no effect of norBNI on distance travelled in either stimulation compartment (Figure 2G; 2-way ANOVA: Drug, F(1,20) = 0.8, *p* = 0.4; Compartment, F(1,20) = 1.8, *p* = 0.2; Drug x Compartment, F(1,20) = 0.1, *p* = 0.7). Administration of norBNI had a significant effect on velocity, but this was not different between stimulation ON and OFF compartments (Figure 2H; 2-way ANOVA: Drug, F(1,20) =7.6, *p* = 0.01; Compartment, F(1,20) = 2.5, *p* = 0.1, Drug x Compartment, F(1,20)= 0.5, *p* = 0.5).

Lastly, in the post-stimulation test session, mice injected with vehicle spent significantly more time in the paired compartment compared to the unpaired compartment, whereas the mice given norBNI showed no significant preference for either compartment (Figure 2E; Vehicle: *t*(6)= 2.7, *p* =0.04; norBNI: *t*(4) = 0.1, *p* =0.2). Taken together, inhibition of KORs did not influence preference for LH orexin/dynorphin optical stimulation in the VTA. However, the ability to recall this preference was inhibited by norBNI.

### Endogenous LH orexin/dynorphin release in the VTA promotes approach to a Pavlovian food cue

To test if optical stimulation of LH orexin/dynorphin in the VTA influences the incentive value of food cues, we used a Pavlovian conditioning procedure where LH orexin/dynorphin inputs were stimulated upon delivery of the conditioned stimulus predicting food pellets (Figure 3A). In the absence of optical stimulation, baseline food value measured by sucrose intake (mCherry: 0.03 ± 0.004 Kcal/g; ChR2 mice: 0.04 ± 0.009 Kcal/g, *t*(16) = 1.9, *p* = 0.07) and baseline cue value measured by lever presses made for the compound light-tone food cue (mCherry: 7.7 ± 2.9 presses; ChR2 mice: 5.4 ± 2.3 presses, *t*(16) = 0.795, *p* = 0.419) were not significantly different between mCherry and ChR2 groups. Next, we examined the number of conditioned responses by quantifying orientations to food cue and magazine approaches during the Pavlovian conditioning sessions across days. Optical stimulation of LH orexin/dynorphin inputs to the VTA of ChR2 mice showed significantly more orienting to the cue compared to mCherry mice (Figure 3B; RM ANOVA: Virus, F(1,12) = 8.7, *p* = 0.02; Day, F(6,72) = 5.5, *p* <0.001; Virus x Day, F(6,72)= 3.5, *p* = 0.004). There was no significant difference between mCherry and ChR2 mice in the number of magazine approaches during the cue (Figure 3C; RM ANOVA: Virus, F(1,12)= 3.4, *p* = 0.09; Day, F(6,72) = 1.9, *p* = 0.09; Day x Virus, F(6,72)= 0.3, *p* = 0.9). We also compared the effect of optical stimulation on entries into the magazine without food cue presentation. Both ChR2 and mCherry mice significantly decreased non-CS magazine entries across conditioning days but this did not differ between mCherry and ChR2 mice (Figure 3D; RM ANOVA: Virus, F(1,13)= 1.3, *p* = 0.3; Day, F(1,9)= 2.2, *p* = 0.006; Virus x Day, F(1,9)= 1.2, *p* = 0.3). These results indicate that both mCherry and ChR2 mice learned the predictive cue value, but optical stimulation of LH orexin/dynorphin inputs to the VTA increased orientation to the food cue.

**Figure 3.**
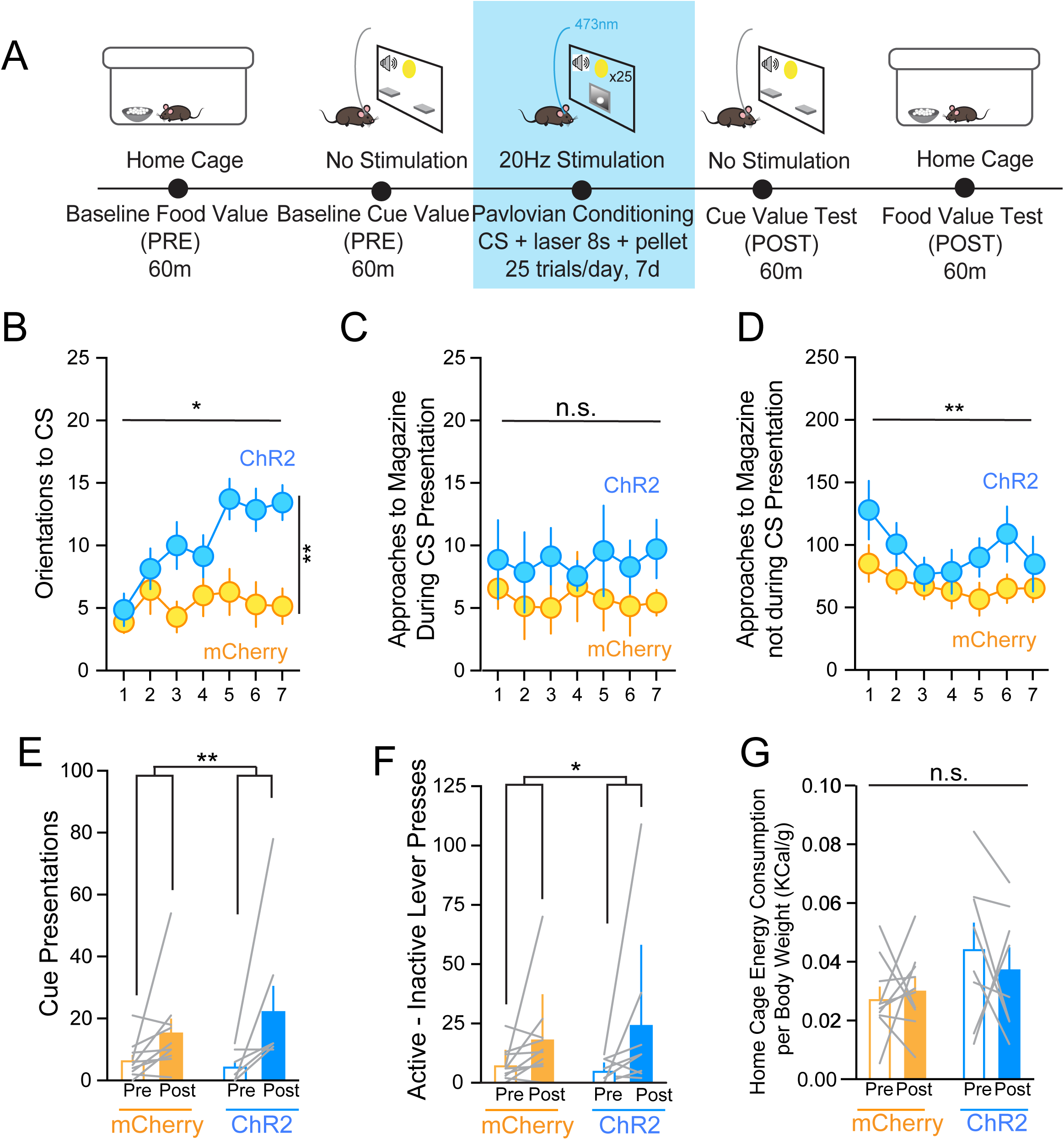
Optogenetic stimulation of LH orexin/dynorphin inputs in the VTA promotes orientations to a Pavlovian food cue. A) Schematic of behavioural paradigm. Blue square denotes intra-VTA LH orexin/dynorphin optogenetic stimulation sessions. B) ChR2 mice (blue) made more orienting responses to the conditioned stimulus (CS) throughout sessions compared to mCherry mice (orange). C) ChR2 and mCherry mice did not differ in the number of magazine approaches made during the CS presentation across conditioning sessions. D) ChR2 and mCherry mice decreased the number of magazine approaches in the absence of cue presentation across conditioning sessions. E) mCherry and ChR2 mice received significantly more cue presentations during the post-conditioning cue value test (filled bars) compared to pre-conditioning baseline cue value assessment (open bars). There was no difference in the number of cue presentations between mCherry and ChR2 mice. F) mCherry and ChR2 mice made significantly more active-inactive lever presses during the post-conditioning cue value test (filled bars) compared to pre-conditioning baseline cue value assessment (open bars). There was no difference in the number of active-inactive responses made for the cue between mCherry and ChR2 mice. G) mCherry and ChR2 mice did not differ in home cage sucrose consumed per body weight at pre-conditioning baseline (open bars) or post-conditioning test (filled bars).

We next assessed whether intra-VTA optical stimulation of LH orexin/dynorphin inputs during Pavlovian conditioning influenced cue value by comparing the number of cue presentations received by lever pressing in the baseline cue value assessment (PRE) and post-conditioning cue value test session (POST). ChR2 and mCherry mice received significantly more cue presentations during the post-conditioning cue value test session compared to pre-conditioning baseline, however, there was no effect of optogenetic stimulation of LH orexin/dynorphin on the number of cue presentations (Figure 3E; RM ANOVA: Virus, F(1,16)= 0.2, *p* = 0.6; Day, F(1,16) = 9.8, *p* = 0.006; Virus x Day, F(1,16) = 1.1, *p* = 0.3). Next, we compared the number of lever presses made for the food cue presentation during the baseline compared to post-conditioning. While both groups of mice significantly increased lever presses made for the cue after conditioning, there was no difference between mCherry or ChR2 mice (Figure 3F; RM ANOVA: Active - inactive lever presses, F(1,16)= 5.4, *p* = 0.03; Virus, F(1,16)= 0.06, *p* = 0.8; Active - inactive lever presses x virus, F(1,16)= 0.4, *p* = 0.5). To determine if the value of the primary reward changed with intra-VTA optical stimulation of LH orexin/dynorphin inputs, we compared home cage baseline sucrose consumed to that consumed during the food value test post-conditioning (Figure 3A,G). Sucrose consumption during the baseline and food value test did not differ significantly between mCherry and ChR2 mice (Figure 3G; RM ANOVA: Time, F(1,16)= 0.1, *p* = 0.7, Virus, F(1,16)= 3.7, *p* = 0.07, Time x Virus, F(1,16)= 0.7, *p* = 0.4). Thus, optical stimulation of LH orexin/dynorphin inputs in the VTA during Pavlovian conditioning did not influence the value of the primary food reward.

### Endogenous LH orexin/dynorphin in the VTA release potentiates electrically evoked mesolimbic dopamine neurotransmission

Given that phasic optical stimulation of VTA dopamine neurons induces RTPP [36] and increased dopamine underlies incentive motivation [1], we next determined whether optical stimulation of LH orexin/dynorphin inputs in the VTA could influence dopamine concentration using in vivo FSCV. Exogenous application of orexin in the VTA increases dopamine release [25], whereas systemic oxR1 antagonists inhibit cocaine-evoked dopamine [24]. However, it is unknown how optogenetic stimulation of LH orexin/dynorphin inputs in the VTA can influence dopamine in the NAc. To examine whether optical stimulation of LH orexin/dynorphin inputs to the VTA could modulate electrically-evoked dopamine (Figure 4A, B), we electrically stimulated dopamine release with or without intra-VTA optical stimulation of LH orexin/dynorphin inputs (Figure 4C-E). Without optical stimulation, electrically-evoked stimulation was not significantly different between mCherry (1.4 ± 0.5 μM) and ChR2 mice (2.3 ± 0.9 μM) (*t*(13) = 1.6, *p* = 0.1).

**Figure 4.**
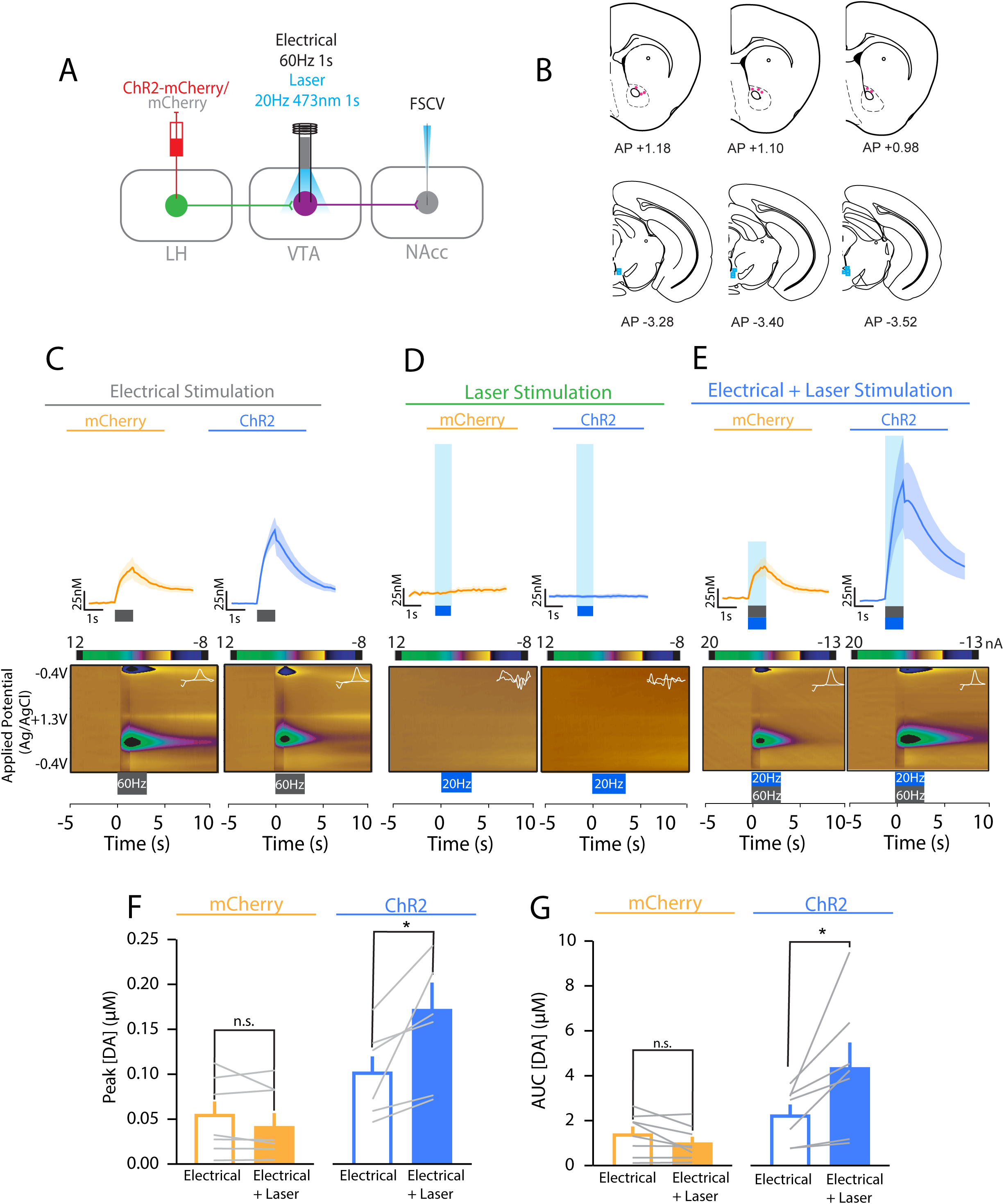
Optogenetic stimulation of LH orexin/dynorphin inputs in the VTA potentiates evoked NAc dopamine release. A)Schematic of viral injection (LH), electrode and optical fibre placement and stimulation parameters (VTA), and FSCV recording location in the NAc core. B)Coronal brain sections (modified from [55]) at 6 different anterior-posterior locations relative to Bregma illustrating the approximate recording (top; pink circles) and stimulating (bottom; blue squares) locations of FSCV data. Approximate electrode tip location was retrospectively determined using the electrode track and recording depths (relative to brain surface). C)Averaged dopamine traces for mCherry (top left, orange) and ChR2 (top right, blue) groups in response to electrical stimulation. Representative false colour plots of dopamine concentration for mCherry (bottom left) and ChR2 (bottom right) mice following electrical stimulation. In all panels, grey box denotes 60Hz 60pulses (1s) electrical stimulation onset and offset. Inset to colour plots, current-voltage plot showing oxidation and reduction peaks for dopamine. Inset, current-voltage plot showing oxidation and reduction peaks for dopamine. D)Averaged dopamine traces for mCherry (top left) and ChR2 (top right) groups in response to laser stimulation. Representative false colour plots of dopamine concentration for mCherry (bottom left) and ChR2 (bottom right) mice following electrical stimulation. In all panels blue box denotes optical stimulation (20Hz) onset and offset. Inset, current-voltage plot showing oxidation and reduction peaks for dopamine. E)Averaged dopamine traces for mCherry (top left) and ChR2 (top right) groups in response to electrical and laser stimulation. Representative false colour plots of dopamine concentration for mCherry (bottom left) and ChR2 (bottom right) mice following electrical and optical stimulation. In all panels blue box denotes optical stimulation (20 Hz) and electrical stimulation (60 Hz) onset and offset. F)There was no significant difference in the peak dopamine concentration ([DA]) recorded for electrical and electrical (open bars) and laser (filled bars) stimulation for mCherry mice (orange). However, ChR2 mice (blue) showed significantly greater peak dopamine in response to electrical and laser stimulation (filled bars) compared to electrical stimulation alone (open bars). G)There was no significant difference in the AUC for dopamine concentration in response to electrical (open bars) and electrical and laser stimulation (filled) for mCherry mice. ChR2 showed a significant potentiation of dopamine AUC in response to electrical and laser stimulation (filled bars) compared to electrical stimulation alone (open bars).

There was a significant effect of virus and optical stimulation on evoked dopamine concentration. While there was no significant potentiation of evoked dopamine peak or area under the curve (AUC) with laser stimulation of LH orexin/dynorphin inputs in the VTA of mCherry mice, peak evoked dopamine concentration (Figure 4F) or AUC (Figure 4G) was significantly potentiated by laser stimulation in ChR2 mice (**Peak:** RM ANOVA: Virus, (F(1,13)= 36.2, *p* = 0.003; Stimulation, F(1,13)= 4.8, *p* = 0.03; Virus x Stimulation, F(1,13) = 10.4, *p* = 0.04; Sidak’s post hoc: mCherry, *p* = 0.2; ChR2, *p* = 0.0001; **AUC:** RM ANOVA: Virus, F(1,13) = 7.8, *p* = 0.02, Stimulation, F(1,13)= 4.5, *p* = 0.04; Virus x Stimulation, F(1,13)= 10.4, *p*= 0.007, Sidak’s post hoc: mCherry, *p* = 0.5; ChR2, *p* = 0.001). We also tested if optical stimulation of intra-VTA of LH orexin/dynorphin inputs alone can evoke dopamine in the NAc core. Optical stimulation did not significantly alter NAc dopamine concentration (AUC) for the 10 s after stimulus offset in either mCherry (0.3 ± 0.2 μM^2^) or ChR2 mice (0.08 ± 0.2 μM^2^) (*t* (12) = 1.363, *p* = 0.1979; Figure 4D). Taken together, intra-VTA optical stimulation of LH orexin/dynorphin inputs alone is insufficient to drive dopamine release in the NAc. However, optogenetic stimulation of LH orexin/dynorphin inputs in the VTA significantly potentiated electrically-evoked dopamine in ChR2, but not mCherry mice.

To examine whether evoked dopamine concentration potentiation by intra-VTA optical stimulation of LH orexin/dynorphin inputs was mediated by oxR1 signalling, we measured the effect of the oxR1 antagonist, SB-334867, on evoked dopamine with or without LH orexin/dynorphin stimulation in the VTA of ChR2 or mCherry mice (Figure 5A-D). Before SB-334867, optical stimulation of evoked dopamine influenced peak concentration (Figure 5Ci; RM ANOVA: Stimulation, F(1,12)= 18.3, *p* = 0.001; Virus, F(1,12) = 0.3, *p* = 0.4, Stimulation x Virus, F(1,12) = 22.1, *p* =0.0005) or AUC (Figure 5Di; RM ANOVA: Stimulation, F(1,12)= 11.7, *p* = 0.005; Virus, F(1,12)= 3.6, *p* = 0.08; Stimulation x virus, F(1,12) = 16.6, *p* =0.002) in the NAc of ChR2 (Sidak’s post hoc: peak: *p* < 0.0001, AUC: *p* = 0.0004), but not mCherry mice (peak: *p* = 0.9, AUC: *p* = 0.8784). After SB-334867, optical stimulation did not significantly alter evoked peak dopamine concentration in the NAc or AUC of either mCherry or ChR2 mice (Figure 5Cii, 4Dii; RM ANOVA: Stimulation, F(1,12) = 1.2, *p* = 0.3; Virus, F(1,12) = 2.4, *p* = 0.2, Stimulation x Virus, F(1,12) = 0.2, *p* = 0.7). Taken together, potentiation of NAc dopamine by stimulation of LH orexin/dynorphin inputs to the VTA was blocked by administration of SB-334867.

**Figure 5.**
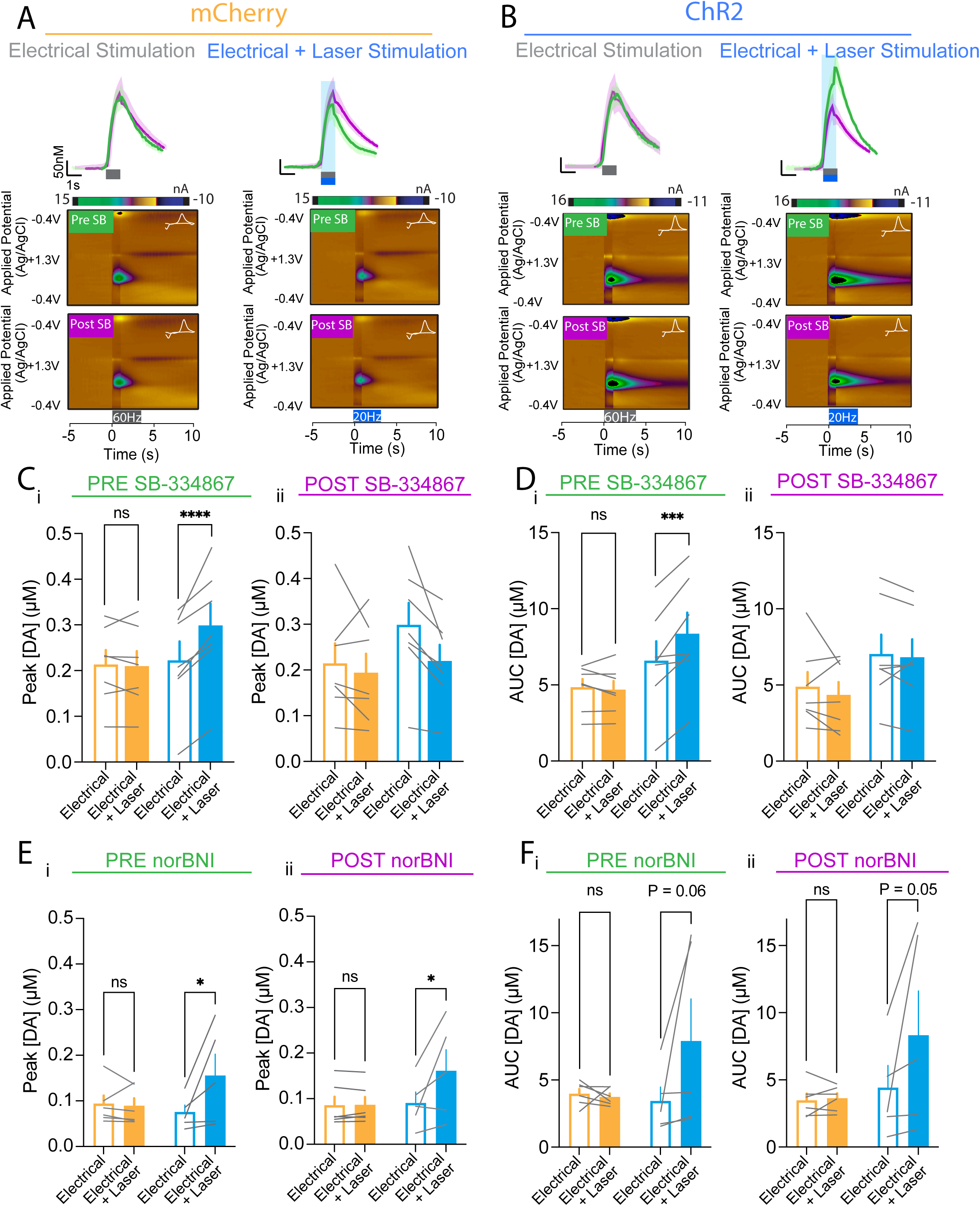
OxR1 is required for potentiation of evoked NAc dopamine release by optogenetic stimulation of LH orexin/dynorphin inputs in the VTA. A) Averaged dopamine traces and representative false colour plots in response to electrical stimulation (top) and electrical and laser stimulation (bottom) for mCherry mice before (green) and after administration of SB-334867 (magenta). Insets to colour plots, current voltage plots showing oxidation and reduction peaks for dopamine. B) Averaged dopamine traces and representative false colour plots in response to electrical stimulation (top) and electrical and laser stimulation (bottom) for ChR2 mice before (green) and after administration of SB-334867 (magenta). Insets to colour plots, current voltage plots showing oxidation and reduction peaks for dopamine. C) (i) Prior to administration of SB-334867, there was no significant difference in the peak dopamine concentration observed in response to electrical (open bars) or electrical and laser stimulation (filled bars) for mCherry mice (orange). For ChR2 mice (blue), laser stimulation (filled bars) significantly potentiated the peak dopamine concentration observed in response to electrical stimulation (open bars). (ii) After administration of SB-334867, there was no significant difference in the peak dopamine concentration observed in response to electrical (open bars) or electrical and laser stimulation (filled bars) for mCherry (orange) or ChR2 mice (blue). D) (i) Prior to administration of SB-334867, there was no significant difference in the AUC of evoked dopamine observed in response to electrical (open bars) or electrical and laser stimulation (filled bars) for mCherry mice (orange). For ChR2 mice (blue), laser stimulation (filled bars) significantly potentiated the AUC observed in response to electrical stimulation (open bars). (ii) After administration of SB-334867, there was no significant difference in the peak dopamine concentration observed in response to electrical (open bars) or electrical and laser stimulation (filled bars) for mCherry (orange) or ChR2 mice (blue). E) (i) Prior to administration of norBNI, there was no significant difference in the peak dopamine concentration observed in response to electrical (open bars) or electrical and laser stimulation (filled bars) for mCherry mice (orange). For ChR2 mice (blue), laser stimulation (filled bars) significantly potentiated the peak dopamine concentration observed in response to electrical stimulation (open bars). (ii) After administration of norBNI, there was a significant difference in the peak dopamine concentration observed in response to electrical (open bars) or electrical and laser stimulation (filled bars) for mCherry (orange) or ChR2 mice (blue). F) (i) Prior to administration of norBNI, there was no significant difference in the AUC of evoked dopamine observed in response to electrical (open bars) or electrical and laser stimulation (filled bars) for mCherry mice (orange) or for ChR2 mice (blue). (ii) After administration of norBNI, there was no significant difference in the peak dopamine concentration observed in response to electrical (open bars) or electrical and laser stimulation (filled bars) for mCherry (orange) or ChR2 mice (blue).

Next, we tested whether evoked dopamine concentration potentiation by intra-VTA optical stimulation of LH orexin/dynorphin inputs was influenced by KOR signalling. We measured the effect of norBNI on evoked dopamine neurotransmission with or without LH orexin/dynorphin stimulation in the VTA of ChR2 or mCherry mice (Figure 5E-G). Before norBNI, optical stimulation of evoked dopamine influenced peak concentration (Figure 5Ei; RM ANOVA: Stimulation, F(1,9)= 4.8, *p* = 0.06; Virus, F(1,9) = 0.5, *p* = 0.5, Stimulation x Virus, F(1,9) = 5.9, *p* =0.04) or AUC (Figure 5Fi: RM ANOVA: Stimulation, F(1,9)= 3.03, *p* = 0.1; Virus, F(1,9)= 1.06, *p* = 0.3; Stimulation x Virus, F(1,9) = 3.8, *p* =0.08) in the NAc of ChR2 (Sidak’s post hoc (a priori test), Peak: *p* = 0.02, AUC: *p* = 0.06), but not mCherry mice (Peak: *p* = 0.9, AUC: *p* = 0.9). After norBNI, optical stimulation significantly increased evoked peak (RM ANOVA: Stimulation, F(1,9) = 5.7, *p* = 0.04; Virus, F(1,9) = 1.2, *p* = 0.3, Stimulation x Virus, F(1,9) = 5.4, *p* = 0.04), but not AUC (RM ANOVA: Stimulation, F(1,9) = 3.5, *p* = 0.08; Virus, F(1,9) = 1.8, *p* = 0.2, Stimulation x Virus, F(1,9) = 3.5, *p* = 0.09) dopamine concentration in the NAc of ChR2 (Sidak’s post hoc (a priori), peak: *p* = 0.0224, AUC: *p* = 0.05), but not mCherry mice (peak: *p* = 1.0, AUC: *p* = 1.0). Taken together, potentiation of NAc peak dopamine by stimulation of LH orexin/dynorphin inputs to the VTA is not significantly altered by administration of norBNI, suggesting that dynorphin does not contribute to LH orexin/dynorphin in the VTA-mediated potentiation of NAc dopamine.

LH orexin/dynorphin neurons are also known to express vesicular glutamate transporter 1 and 2 [35] and optical stimulation of these neurons in the LH can evoke AMPA excitatory postsynaptic currents (EPSCs) onto histaminergic neurons [37,38]. Therefore, we wanted to determine whether optical stimulation of LH orexin/dynorphin inputs in the VTA could evoke AMPA EPSCs. However, we found that intra-VTA stimulation of LH orexin/dynorphin did not evoke AMPA EPSCs recorded at -70 mV (data not shown). Given that exogenous application of orexin can potentiate electrically evoked NMDA currents onto VTA dopamine neurons [26,39], we next tested if intra-VTA stimulation of LH orexin/dynorphin inputs could potentiate NMDA currents of dopamine neurons. Optical stimulation of LH orexin/dynorphin inputs potentiated electrically evoked NMDA currents onto dopamine neurons (Supplemental Figure 2A-C; Wilcoxon matched-pairs signed rank test, *p* = 0.01, W = 62, N/n = 12/6). However, pre-application of SB-334867 blocked the potentiation of NMDA currents by optical stimulation and induced a depression of evoked NMDA EPSCs (Supplemental Figure 2D-F; Wilcoxon matched-pairs signed rank test, *p* = 0.03, W = -21, N/n = 6/4). Application of norBNI also blocked potentiation of NMDA currents (Supplemental Figure 2G-I, Wilcoxon matched-pairs signed rank test, *p* = 0.9, W = -31, N/n = 11/5). Taken together, while optical stimulation of LH orexin/dynorphin inputs in the VTA does not produce an excitatory synaptic current onto dopamine neurons alone, it can potentiate existing excitatory synaptic currents via a neuromodulatory effect.

## DISCUSSION

Here, we found that endogenous LH orexin/dynorphin release in the VTA significantly potentiates evoked mesolimbic dopamine neurotransmission, an effect dependent on oxR1, but not KOR signaling. Furthermore, activation of LH orexin/dynorphin in the VTA can produce place preference that is also observed after cessation of optical stimulation, consistent with previous work demonstrating that systemic SB-334867 can inhibit the expression of conditioned place preference (CPP) for food[40] or drugs of abuse[14,23]. During optical stimulation of LH orexin/dynorphin in the VTA, orientations to a food cue is enhanced, a key motivational feature of reward cues. Finally, while optical stimulation of LH orexin/dynorphin into the VTA alone did not produce EPSCs, it potentiated evoked NMDA EPSCs, suggesting a neuromodulatory role in driving these behaviours. Thus, LH orexin/dynorphin in the VTA may promote motivated reward-seeking behaviours through potentiating excitatory synaptic transmission in the VTA and NAc dopamine.

Our results extend previous work demonstrating that exogenously applied orexin in the VTA increases striatal dopamine. Inhibition of oxR1 in the VTA disrupts cocaine-potentiated evoked phasic dopamine release measured with FSCV in the rat caudate putamen [24]. Similarly, intra-VTA orexin application augments dopamine [41] and potentiates cocaine-induced tonic dopamine release in rat NAc core measured with microdialysis [25]. Intra-VTA optical stimulation of LH orexin/dynorphin inputs alone was not sufficient to augment NAc dopamine but required phasic activation of VTA dopamine neurons to increase dopamine in the NAc core. Consistent with this, intra-VTA orexin can increase phasically-evoked dopamine release in the caudate putamen with or without intravenous cocaine [25]. Intra-VTA KOR activation can suppress NAc dopamine and aversion learning [42] and that intra-VTA blockade of KORs disrupts decreased dopamine associated with aversive stimuli [30], suggesting that dynorphin released during aversive stimuli [43] decreases NAc dopamine through activation of KORs in the VTA [30]. However, inhibition of KORs during LH orexin/dynorphin optical stimulation did not alter phasically-evoked dopamine responses, suggesting that it is unlikely that the source of VTA dynorphin released with aversive stimuli is from LH orexin/dynorphin neurons. Consistent with this, intra-VTA administration of norBNI blocked dopamine release in the medial prefrontal cortex but not the NAC [44,45]. Taken together, potentiation of phasic dopamine is driven by LH orexin rather than LH dynorphin during optical stimulation of the LH orexin/dynorphin input to the VTA.

Glutamate is colocalized with LH orexin/dynorphin neurons [35] and could be released into the VTA with optical stimulation, leading to enhanced dopamine release [46]. We found that optical stimulation of LH orexin/dynorphin inputs to the VTA did not produce an EPSC in the VTA, consistent with few synaptic contacts of LH orexin/dynorphin neurons onto dopamine neurons [18]. However, optical stimulation of LH orexin/dynorphin neurons potentiated evoked NMDA responses, suggesting a neuromodulatory effect of these peptides on excitatory synaptic transmission onto VTA dopamine neurons. Indeed, exogenously applied orexin in the VTA increases NMDA currents and promotes synaptic plasticity of dopamine neurons [39]. As such, LH orexin/dynorphin may increase the gain of excitatory inputs onto VTA dopamine neurons. Given that NMDA receptor activation in the VTA is required for phasic dopamine [46,47], LH orexin/dynorphin in the VTA may require coincident NMDA receptor activation, which likely occurs with 60 Hz stimulation, to potentiate NAc dopamine and thus impact reward processes dependent on mesolimbic dopamine. Potentiation of NMDA was blocked by either oxR1 or KOR antagonists, suggesting that orexin and dynorphin are required. One possibility is that co-release of orexin and dynorphin induce NMDA receptor potentiation via both pre- and postsynaptic mechanisms. Activation of postsynaptic oxR1 on dopamine neurons can increase trafficking of NMDA receptors to the synapse [39] as well as endocannabinoid-mediated suppression of GABAergic inputs leading to a shift in strength of excitatory synapses [48]. In addition to KOR suppression of EPSCs [49], KOR activation can supress GABAergic synaptic transmission onto dopamine neurons [9] and prevent synaptic strengthening of GABAergic inputs to dopamine neurons [50]. Thus, optical stimulation of orexin and dynorphin could potentiate NMDA EPSCs via concerted oxR1 and KOR action leading to a shift in inhibitory to excitatory balance, but this speculation requires further study.

Optical stimulation of LH orexin/dynorphin inputs in the VTA produced a strong place preference for a context paired with that stimulation. Continuous pairing of the intra-VTA LH orexin/dynorphin stimulation transformed a neutral stimulus (compartment) to a conditioned stimulus, whereby mice spent more time in the paired compartment. LH orexin/dynorphin stimulation in the VTA induced a progressive increase in preference for the paired compartment over the 3 pairing days in an oxR1-dependent manner, suggesting that orexin action in the VTA plays a role in acquisition of this preference. Notably, SB-334867 decreased general locomotor activity. However, this did not affect exploration of either compartment as mice explored both arenas equally, suggesting that the general effects of SB-334867 on locomotor activity did not impair the ability of mice to make an association with a rewarding compartment. Inhibition of oxR1 in the VTA increases reward thresholds for electrical self-stimulation and decreases impulsive responding on a 5-choice serial reaction time task and cocaine self-administration[21]. However, these effects are blocked by pretreatment with a KOR antagonist [21], suggesting that orexin in the VTA can negate the aversive effects of dynorphin. We tested if optically stimulated dynorphin release contributed to RTPP using the KOR inhibitor, norBNI. Pretreatment with a KOR agonist in the VTA potentiates preference for cocaine-associated contexts in a CPP paradigm [51,52] and cocaine-evoked NAc dopamine [52]. However, here, norBNI did not influence the preference for optical stimulation of VTA LH orexin/dynorphin inputs or potentiation of evoked dopamine release. Taken together, rather than LH dynorphin acting presynaptically to disinhibit NAc core-projecting dopamine neurons, similar to cocaine, leading to potentiation of place preference or dopamine release, it is likely that LH orexin/dynorphin release may be modulating different circuits, with the effects of LH orexin predominating in the VTA-NAc projection[10]. Thus, optical stimulation of LH orexin/dynorphin in the VTA is sufficient to produce a contextual preference in the absence of any tangible primary reward, suggesting that LH orexin/dynorphin stimulation in the VTA is critical for underlying reward-seeking behaviours.

24 hours after cessation of optical stimulation of LH orexin/dynorphin inputs in the VTA, a continued place preference was observed. This suggests a long-term encoding of a preference for the environment paired with endogenous orexin release, rather than a transient behavioural influence of the stimulation itself. These data uphold previous findings that intra-VTA oxR1 signaling attenuates CPP to morphine or cocaine[23,41], suggesting that LH orexin/dynorphin in the VTA is sufficient for associating the context with both natural rewards and drugs of abuse. However, when KORs were blocked during the acquisition of the RTPP, the post-stimulation preference test was no longer significantly different from controls. Further research should test whether LH dynorphin release during the development of preference for intra-VTA LH orexin/dynorphin stimulation is required for the formation of this contextual association.

One possibility for differential effects of LH orexin vs. dynorphin on RTPP and dopamine release may be due to different optical release requirements of orexin compared to dynorphin. For example, while 20 Hz optogenetic stimulation released orexin somatodendritically [37,38] or at locus coeruleus terminals [53], dynorphin can be released at 5 Hz within the bed nucleus of the stria terminalis [54]. Peptide release requirements could vary from region-to-region based on the number of dense core vesicles. Immuno-electron microscopy work has indicated that orexin and dynorphin are co-expressed in the same dense core vesicles within the LH [21] and presumably would be released at sites in the VTA under the same release requirement. Thus, we speculate that the difference in LH orexin vs dynorphin effects on RTPP and dopamine release are due to differential oxR1 and KOR expression on VTA originating circuits, rather than differences in release requirements.

Because LH orexin/dynorphin in the VTA was necessary and sufficient for associating the context with preference for stimulation, we assessed if LH orexin/dynorphin in the VTA was involved in motivation towards a food reward and/or a food cue. Optical stimulation of LH orexin/dynorphin inputs in the VTA increased orientation to a food cue, suggesting that endogenous orexin release in the VTA modulates the ability of food cues to direct reward-seeking behaviour. However, we did not observe any influence of prior LH orexin/dynorphin stimulation in the VTA on subsequent conditioned reinforcement behaviour, suggesting no enduring changes in the incentive value of food cues in this paradigm. Furthermore, it is important to note that the conditioning effect we observed may be to the rewarding features of LH orexin/dynorphin stimulation itself, rather than the sucrose per se. We did not observe changes in the incentive value of the primary reward, suggesting that LH orexin/dynorphin in the VTA is only implicated in the motivational processes relating to conditioned stimuli.

In conclusion, this study demonstrates a significant role for endogenous orexin release in the VTA in the development of RTPP, food cue-directed motivation, and potentiation of excitatory synaptic transmission and mesolimbic dopamine neurotransmission. Given that optical stimulation of LH orexin/dynorphin terminals in the VTA likely release dynorphin in addition to orexin, our results demonstrate that the effects of orexin predominate in mediating reward-seeking behaviour. These findings shift forward our understanding of neural circuitry underpinning reward-related processes and highlight that neuromodulation in distinct target regions may have independent or opposing effects. These findings not only contribute to our knowledge of general reward processes but, alongside previous work demonstrating a role for orexin in addiction, add support for targeting orexinergic systems as a possible treatment of addiction.

## Data accessibility

Data will be made available upon request.

## Acknowledgements

The authors would like to acknowledge the Hotchkiss Brain Institute optogenetic core facility and the advanced microscopy facility for their technical support.

## Author Contributions

C.S.T. designed, performed and analyzed all experiments with supervision of S.L.B. M.R. and A.M. performed and analyzed behavioural experiments with supervision of S.L.B and C.S.T. M.Q. performed immunohistochemistry experiments. A.M. performed the electrophysiological experiments in the VTA. C.B. performed electrophysiological recordings in the LH. C.S.T. and S.L.B. wrote the manuscript and agree to be accountable for all aspects of the work.

