## Supplemental for "Optogenetic stimulation of lateral hypothalamic orexin/dynorphin inputs in the ventral tegmental area potentiates mesolimbic dopamine neurotransmission and promotes reward-seeking behaviours"

**SUPPLEMENTAL MATERIALS**

**Extended Methods**

**Location**

This research was performed at the University of Calgary which is located on the unceded traditional territories of the people of the Treaty 7 region in Southern Alberta, which includes the Blackfoot Confederacy (including the Siksika, Piikuni, Kainai First Nations), the Tsuut'ina, and the Stoney Nakoda (including the Chiniki, Bearspaw, and Wesley First Nations). The City of Calgary is also home to Metis Nation of Alberta, Region III.

**Subjects**

Adult male and female orexin-EGFP-2A-cre mice (post-natal day 60-90) (originally from the Yamanaka lab at the Nagoya University [1], bred locally) were group-housed (3-5 per cage) with *ad libitum* access to water and food (temperature ($21 \pm 2$ºC), humidity-controlled (30-40%), 12h reverse light/dark cycle). All testing was conducted during the active (dark) phase of the cycle. All experimental procedures adhered to ethical guidelines established by the Canadian Council for Animal Care and animal use protocols approved by the University of Calgary Animal Care and Use Committee.

**Viral Infusion and Optical Fibre Implantation Surgeries**

All mice received bilateral infusions of either channelrhodopsin ('ChR2') (AAV2/8-EF1a-DIO-hChR2(H134R)-mCherry; Neurophotonics, Centre de Recherche CERVO, Laval, QC) or control ('mCherry') (AAV2/8-hSyn-DIO-mCherry; Neurophotonics) virus. Some orexin-cre mice (in Supplemental Figure 1) were injected with rAAV2-EF1a-DIO-hCHR2(H134R)-eYFP. Mice were anaesthetised with isofluorane gas and secured in a stereotaxic frame (David Kopf Instruments, Tujunga, CA). All stereotaxic measurements were made relative to bregma. Viral injections were performed using a microinjector (Nano-inject II; Drummond Scientific Company, Broomall, PA, USA). Each mouse received 6 infusions into the LH (100 nl per infusion; 23.1 nl/s), 3 in each hemisphere (anterior-posterior (AP) -1.35; mediolateral (ML) $\pm$ 0.9; dorsoventral (DV) -5.2, -5.1, -5.0) for a total of 300 nl per hemisphere. After each infusion, the microinjector was left in place for 3 min to allow diffusion of virus away from the needle tip. After all infusions were complete in one hemisphere, the microinjector was left in place for an addition 5 min, 500 $\mu$m dorsal of the final injection location to allow diffusion of the virus through the brain tissue. All mice received pre- and postoperative analgesic (Ketoprofen 5 mg/kg or Meloxicam 3 mg/kg, subcutaneous) and were returned to their home cages and allowed to recover for 6-8 weeks prior to further experimental procedures. The location of the virus expression was examined post hoc.

Mice included in behavioural experiments received a second surgery to implant optical fibres targeted at the VTA, 6 weeks after viral infusions. Again, mice were anaesthetised with isofluorane and secured in a stereotaxic frame (as above). Bilateral optical fibres (Doric Lenses Inc; Quebec City, QC, Canada) were lowered to just dorsal of the VTA (AP -3.5; ML $\pm$ 0.5; DV -3.8, -4.0) and secured with Metabond (C & B Metabond, Parkell Inc, Edgewood, NY, USA) and dental cement (Lang Dental Ltd, Rochester, NJ, USA). Mice were returned to their home cage and allowed to recover for 1 week prior to behavioural testing.

**Real-Time Place Preference (RTPP)**

Orexin-cre ChR2 (n=8) and Orexin-cre mCherry (n=10) mice implanted with bilateral optical fibres underwent 3 stages of real-time place preference paradigm:

1. *Baseline Session (day 1).* Mice were attached to an extended patch cable and placed in the real-time place preference arena (50cm x 25cm x 50cm; underlit black Plexiglas, [2]) for 20 min without receiving any stimulation, to record any innate preference for either compartment.
2. *Stimulation Sessions (days 2-4).* Mice were attached to an extended patch cable connected to a laser (473nm, 20Hz, 5mW, 5ms pulses) and placed into the arena again. One compartment of the arena was paired with laser activation (stimulation ON), whereas one compartment was paired with no stimulation (stimulation OFF). The arena compartment paired with stimulation was counter-balanced across mice. Each session lasted 60 min. Mice were tested at the same time every day.
3. *Test Session (day 5).* Mice were again attached to the extended patch cable and allowed to explore the arena for 20 min without receiving any stimulation.

Data were collected and laser stimulation (473 nm, 20 Hz, 5 mW, 5 ms pulses, 1 s duration) triggered using EthoVision XT software (Noldus Information Technology, Wageningen, Netherlands).

For the experiment involving the OXR1 antagonist, SB-334867 or the KOR antagonist, norBNI, male and female orexin-cre mice were infused with ChR2 (n = 10 for SB-334867 and n = 14 for norBNI) in LH and implanted with optical fibres targeted at the VTA. Mice underwent the RTPP as above. However, prior to each conditioning session (day 2-4), all mice received an intraperitoneal injection of either SB-334867 (n = 5, 15 mg/kg) (HelloBio Inc., Princeton, NJ, USA; dissolved in vehicle 10% hydroxypropyl-beta-cyclodextrin (HPBCD) and 2% dimethyl sulfoxide (DMSO) in sterile water w/v), or vehicle (n = 5), ), or norBNI (n = 7, 15 mg/kg) or vehicle (n = 7) 15 min prior to being placed into the arena.

**Pavlovian Conditioning**

Orexin-cre ChR2 (n=8) or mCherry (n=10) mice implanted with bilateral optical fibres target at the VTA were used for both RTPP and Pavlovian conditioning in a counter-balanced order.

*Baseline Food Value (2 days).* Mice received ~20 sucrose pellets (Bio-Serv, Flemington, NJ, USA) in their home cage during the active phase of their circadian cycle to overcome neophobia. Next, mice were individually placed into a fresh home cage with a small amount of their bedding in the experimentation room and given access to sucrose pellets for one hour for two days. The amount of sucrose consumed was recorded for each day and compared between groups.

*Baseline Cue Value (3 days).* Mice were next attached to an optical fibre patch cable and placed into a sound-attenuated mouse behavioural conditioning chamber (Med Associates, St. Albans, VT, USA; dimensions: 15.24 cm x 13.34 cm x 12.7 cm). The chamber was equipped with a house light, 2 retractable levers, one cue light, and one sonalert (60-65 dB). Mice were allowed to explore the chamber and make responses on the two levers for 1 h for 3 consecutive days, with session duration signalled by house-light illumination. Responses on one lever (‘active’; counterbalanced across subjects), resulted in the presentation of a light and tone for 8 s. Responses on the other lever (‘inactive’) was inconsequential but recorded.  Active – Inactive responses on day 3 of baseline cue value assessment was compared between groups.

*Pavlovian Conditioning (7 days).* Behavioural chambers were equipped with a magazine, allowing delivery and retrieval of sucrose pellets. Mice were habituated to the operant conditioning chambers with the levers retracted for 1-3 days prior to Pavlovian conditioning. During this period, mice retrieved sucrose pellets from the magazine in session where 25 pellets were delivered to the magazine on a variable time schedule (30 s; inter-trial interval [ITI] 10-60 s). Mice remained in the chamber for 1 h to maximize familiarity with sucrose pellets and environment. Sessions were repeated until mice consumed at least 15 sucrose pellets in one session. Next, mice underwent daily sessions of Pavlovian Conditioning for 7 consecutive days. Mice were connected to an extended patch cable and laser for each session. Each session consisted of 25 trials in which one sucrose pellet was delivered into the magazine, the cue light was illuminated and the sonalert sounded for 8 s. Laser stimulation (as above) lasted for the 8 s duration of the cue presentation for all mice. All trials were paired (sucrose delivery and cue onset at the same time) on a variable time schedule averaging 90 s, with ITI of 30-150 s. Each session lasted approximately 40 min. Duration of session was signalled by illumination of the house-light.

*Cue Value Test (1 day).* 24 h after the last Pavlovian conditioning session, mice were returned to the behavioural chamber. The chamber was equipped with 2 levers (one active; one inactive in locations matching cue baseline testing; food magazine removed). As in cue baseline sessions, mice were given 1 h to respond for presentation of the compound light-tone cue. Responses were compared to each individual mouse's responses in the last session of cue baseline testing.

*Food Value Test (1 day).* As in pre-conditioning food value baseline assessment, mice were placed into a novel home cage with ~20 sucrose pellets in the experimentation room for one hour. The weight of sucrose pellets consumed was recorded and compared to the last day of food baseline testing.

*Video Analysis.* All Pavlovian conditioning sessions were video recorded. The videos were then scored for approach and orientation (including latency to orient and latency to approach) to the cue by experimenters who were blinded to experimental conditions. Videos were each scored twice to provide inter-rater reliability.

##### ***In vivo* Fast-Scan Cyclic Voltammetry (FSCV)**

Mice (ChR2 n= 7; mCherry n=7) were anaesthetized with intraperitoneal injection of 25% urethane (1.0-1.5 g/kg) dissolved in sterile saline. Once deeply anaesthetised, mice were secured in a stereotaxic frame inside a Faraday cage. Small craniotomies were then made above the NAc (AP +1.0; ML +1.0) and the VTA (AP -3.5; ML +0.5) and contralateral cortex (AP +1.8; ML -2.0). A chlorinated silver (Ag/AgCl) reference electrode was implanted in the contralateral cortex (DV -0.8) and cemented in place (C&B metabond, as above).

A recording electrode (details in [3]; Goodfellow, Coraopolis, PA, USA) was lowered to the right NAc core (~DV -3.8) as previously described [4]. FSCV data were collected using TarHeel CV (ESA Biosciences Inc., Chelmsford, MA, USA) and High-Definition Cyclic Voltammetry Suite (University of North Carolina, Chapel Hill). A bipolar stimulating electrode (P1 Technologies [formerly Plastics One], Roanoke, VA, USA), combined with an optical fibre (constructed in house, ~30 μM shorter than the electrode) was incrementally lowered, in 100 μm steps, into the VTA.

Experimental data were collected in 120 s files (stimulation onset 5 s into the recording). 10 recordings of electrical stimulation (60 Hz, 60 pulses; 120 μA), 10 recordings of optical stimulation (laser as above; 473 nm, 20 Hz, 20 mW, 5 ms pulses, 1s duration), and 10 recordings of combined electrical (as above) and optical stimulation (as above) were made in a counter-balanced order across subjects.

Carbon fibre microelectrodes were pre-calibrated to known concentrations of dopamine (250 nM and 1000 nM) as previously described [5] to allow conversion of recorded *in vivo* signals to changes in dopamine concentration using chemometric analysis, principal component regression, and residual analyses using TarHeel and HDCV (as above; [6]). Training set data were established from collating numerous electrochemical signatures of dopamine at various electrical stimulation parameters compared to several pre-established dopamine concentrations. Data were analysed to calculate the peak dopamine and area under the curve for the 10 s period following stimulation onset. Data were compared between electrical, laser, and combined electrical-laser stimulation to determine the effect of LH orexin/dynorphin intra-VTA stimulation on mesolimbic dopamine neurotransmission.

For SB-334867 (ChR2 n = 7; mCherry n = 7) or norBNI (ChR2 n = 6; mCherry n = 5) injections, orexin-cre mice infused with either ChR2 or mCherry virus (as above). We conducted anaesthetised FSCV data acquisition as above, followed by an intraperitoneal injection of SB-334867 (15 mg/kg) or norBNI (15 mg/kg). After 15 min, we recorded evoked responses to electrical stimulation and to electrical + optical stimulation and compared the area under the curve and peak dopamine concentration to recordings made prior to administration of the antagonist.

**Electrophysiology**

**Slice preparation:** All electrophysiological recordings were performed in slice preparations from adult male and female orexin-cre mice (at least 4 months old) that were injected with either rAAV2-EF1a-DIO-hCHR2(H134R)-eYFP (firing experiments, Supplemental Figure 1) or AAV2/8-hSyn-DIO-hChR2(H134R)-mCherry (EPSC experiments, Figure 5) into the LH (as above) 6 weeks prior to patch clamp recordings. Mice were deeply anaesthetized with isoflurane and intracardially perfused with N-methyl D-glucamine (NMDG) solution of composition (in mM): 93 NMDG, 2.5 KCl, 1.2 $NaH_2PO_4.H_2O$, 30 $NaHCO_3$, 20 HEPES, 25 D-glucose, 5 sodium ascorbate, 3 sodium pyruvate, 2 thiourea, 10 $MgSO_4.7H_2O$, 0.5 $CaCl_2.2H_2O$ and saturated with 95% O2–5% $CO_2$. Mice were then decapitated and horizontal midbrain sections (250 µm) containing the VTA were cut in NMDG solution using a vibratome (VT1200, Leica Microsystems, Nussloch, Germany). Slices were recovered in warm NMDG solution (32 °C) saturated with 95% O2–5% $CO_2$ for 10 min before being transferred to a holding chamber containing artificial cerebrospinal fluid (ACSF) of composition (in mM): 126 NaCl, 1.6 KCl, 1.1 $NaH_2PO_4$, 1.4 $MgCl_2$, 2.4 $CaCl_2$, 26

$NaHCO_3$, 11 glucose (32 °C); equilibrated with 95% O2 / 5% $CO_2$ for at least 45 min before recording.

**Electrophysiology:** Slices were transferred to a recording chamber on an upright microscope (Olympus BX51WI) and continuously superfused with ACSF (2 $mL.min^{-1}$, 34 °C). LH orexin/dynorphin neurons were identified by green fluorescence. Putative VTA dopamine neurons were visualized with a 40X water immersion objective using Dodt gradient contrast optics. Whole-cell voltage-clamp recordings (holding potential = -70 mV) of synaptic currents were made using a MultiClamp 700B amplifier (Axon Instruments, Molecular Devices). VTA dopamine neurons were identified by morphological and electrophysiological characteristics (fusiform shape, capacitance > 50 pF, presence of H-current (Ih)). These neurons were further identified as dopamine neurons with post-hoc immunohistochemistry for tyrosine hydroxylase. Recording electrodes (3-5 MΩ) for measuring firing rates in current clamp were filled with potassium-D-gluconate internal solution (in mM): 136 potassium-D-gluconate, 4 MgCl2, 1.1 HEPES, 5, EGTA, 10 sodium creatine phosphate, 3.4 Mg-ATP and 0.1 Na2GTP and 0.2% biocytin. After breaking into the cell, Ih currents were recorded in voltage-clamp mode using a voltage step to -130 mV to dopamine neurons voltage-clamped at -70 mV. Ih was determined as the change in current between ~30 ms and 248 ms after the voltage step was applied.

Because most dopamine neurons ceased firing within 5 minutes of recording, current-step induced firing was used to assess the effects of different frequency stimulation of VTA dopamine neuronal firing. For current-step experiments, the membrane potential for each neuron was set to -60 mV by DC injection via the patch amplifier and a series of 5 current pulses

(250 ms in duration, 5-25 pA apart, adjusted for each cell) were applied every 45 seconds, where the minimum current amplitude was set for each cell so that the first pulse was subthreshold and did not yield firing. We optogenetically stimulated LH inputs in the VTA over a range of frequencies (5, 20 or 30 Hz for 10 s) from a light-emitting diode (LED) blue light source (470nm) directly delivered the light path through the Olympus 40X water immersion lens.

For evoked excitatory postsynaptic currents (EPSCs), a bipolar tungsten-stimulating electrode was placed 100–300 µm rostral to the cell being recorded and stimulated at 0.1 Hz. Recording electrodes (3-5 MΩ) for measuring EPSCs were filled with CeMeSO3 internal solution containing (in mM): 117 CeMeSO3, 2.8 NaCl, 20 HEPES, 0.4 EGTA, 5 TEA, 5 MgATP and 0.5 NaGTP and 0.2% biocytin. AMPA EPSCs were recorded at -70 mV in the presence of picrotoxin (100 µM). NMDA EPSCs were evoked by electrically stimulating afferents while dopamine neurons were voltage clamped at +40 mV in the presence of 100 µM picrotoxin and 10 µM DNQX to block $GABA_A$ and AMPA receptors respectively. NMDAR traces were constructed by averaging EPSCs elicited at +40 mV. EPSCs were filtered at 2 kHz, digitized at 5–10 kHz and collected online. After obtaining a minimum 5 min stable baseline of evoked NMDA EPSCs, we optogenetically stimulated LH inputs in the VTA at 20 Hz for 10 s from a light-emitting diode (LED) blue light source (470nm) directly delivered the light path through the Olympus 40X water immersion lens. SB334867 and NorBNI (Tocris) stock solutions were both dissolved in 100% DMSO and were bath applied at 1µM in 0.01% DMSO throughout the experiment. Data were averaged in 5 min bins, normalized to baseline, which is defined as the average EPSC amplitude of 5 min before stimulation, and presented as average across cells ± SEM.

**Histology**

For behavioural experiments, to check viral transfection, optical fibre placement, and colocalization of orexin and mCherry virus, mice were deeply anesthetized with isoflurane and transcardially perfused with phosphate buffered saline (PBS) and then with 4% paraformaldehyde (PFA). Brains were dissected and post-fixed in 4% PFA at 4°C overnight, then switched to 30% sucrose. Coronal frozen sections were cut at 30 µm using a cryostat. Sections were heated in citrate buffer (pH=6.0) at 80-85°C for 40 min to expose antigen. 10% goat and donkey serums were applied to block non-specific binding for 1 h. Sections were then incubated with primary antibody rabbit anti-orexin 1:500 (Phoenix pharmaceuticals, H-003-30) and chicken red fluorescent protein (RFP) 1:2000 (Rockland, 600-901-379) in 1% BSA for 48 h at room temperature followed by incubation with secondary antibody Alexa Fluor 488 donkey anti-rabbit and Alexa Fluor 594 goat anti chicken 1:400 for 1 h on LH sections and for 4-5 h on VTA sections.

For FSCV experiments, to check for electrode placements and viral transfection, mice were transcardially perfused with PBS and then 4% PFA. Brains were dissected and post-fixed in 4% PFA at 4°C overnight, then switched to 30% sucrose. Coronal frozen sections were cut at 30 µm using a cryostat. Sections were blocked in 10% donkey serum for 1 h before incubation in primary antibody rabbit anti-RFP (Rockland, 600-401-379) 1:2000 in 1% BSA for 1 h followed by incubation with secondary antibody Alexa Fluor 594 donkey anti rabbit 1:400 for 1 h.

For electrophysiology experiments, slices were fixed in 4% PFA overnight at 4$^{o}$C and then rinsed in PBS, blocked in 10% normal goat serum, incubated with monoclonal mouse anti-TH antibody (Sigma, 1:1000, T1299) at room temperature 24 hours followed by incubation with Alexa Flour 488 goat anti mouse (1:400) for 4 hours at room temperature. To co-label biocytin,

Dylight 594 streptavidin (Sigma, 016-510-084, 1:200) was applied and incubated for 2 hours at room temperature.

For all experiments, slices were mounted with Fluoshield (Sigma). All images were obtained on an Olympus virtual slide microscopy VS120-L100-W with a 10x objective (Olympus Canada Inc., Ontario, Canada) and a Leica confocal microscopy TCS SP8 with a 25x objective (Leica Microsystems Inc., Ontario, Canada).

**Statistics**

All statistical analyses were completed using GraphPad Prism 8 or 9. All values are expressed as mean ± standard error of the mean (SEM). The alpha risk for the rejection of the null hypothesis was set to 0.05. All data met criteria for normality unless otherwise specified. All post hoc tests were conducted with Sidak's correction for multiple comparisons, unless otherwise stated.

**Supplemental Figure 1. Optogenetic manipulation of LH orexin/dynorphin neurons.**

A) (left) Patch clamp recording of an rAAV2-EF1a-DIO-hCHR2(H134R)-eYFP expressing LH orexin/dynorphin neuron from an orexin-cre mouse. (right) 5x image of the recording pipette in the LH from a horizontal section of an Orexin-cre mouse.

B) Photocurrent induced by increasing flash duration (0 to 100 ms in 10 ms increments.

C) The response of LH orexin/dynorphin neurons to increasing pulse trains (5-30 Hz) in voltage-clamp recording conditions.

D) The response of LH orexin/dynorphin neurons to increasing pulse trains (5-30 Hz) in current-clamp recording conditions

E) Schematic demonstrating electrophysiological experiment testing the effects of optical stimulation of LH orexin/dynorphin inputs in the VTA on dopamine neuronal firing.

F) Time course of evoked firing (100 pA step) of dopamine neurons before and after 5 Hz, 20 Hz, or 30 Hz (10 s) stimulation.

**Supplemental Figure 2. Optogenetic stimulation of LH orexin/dynorphin inputs to VTA neurons can potentiate NMDA currents.**

A) Time course of electrically evoked NMDA ESPCs of VTA dopamine neurons before and after 20Hz, 10 s optical stimulation (green line) of LH orexin/dynorphin inputs.

B) Bar graph of averages and individual evoked NMDA responses before (open) and 5 min after (filled) optical stimulation of LH orexin/dynorphin inputs to VTA dopamine neurons.

C) Example traces evoked at +40 mV before (black) and after (blue) optical stimulation of LH orexin/dynorphin inputs to VTA dopamine neurons.

D) Time course of electrically evoked NMDA ESPCs of VTA dopamine neurons before and after 20Hz, 10 s optical stimulation (green line) of LH orexin/dynorphin inputs in the presence of SB-334867 (1 μM) throughout the experiment.

E) Bar graph of averages and individual evoked NMDA responses before (open) and 5 min after (filled) optical stimulation of LH orexin/dynorphin inputs to VTA dopamine neuron in the presence of SB-334867 (1 μM).

F) Example traces evoked at +40 mV before (black) and after (blue) optical stimulation of LH orexin/dynorphin inputs to VTA dopamine neurons in the presence of SB-334867 (1 μM).

G) Time course of electrically evoked NMDA ESPCs of VTA dopamine neurons before and after 20Hz, 10 s optical stimulation (green line) of LH orexin/dynorphin inputs in the presence of norBNI (1 μM) throughout the experiment.

H) Bar graph of averages and individual evoked NMDA responses before (open) and 5 min after (filled) optical stimulation of LH orexin/dynorphin inputs to VTA dopamine neuron in the presence of norBNI (1 μM).

I) Example traces evoked at +40 mV before (black) and after (blue) optical stimulation of LH orexin/dynorphin inputs to VTA dopamine neurons in the presence of SB-334867 (1 μM) and norBNI (1 μM).

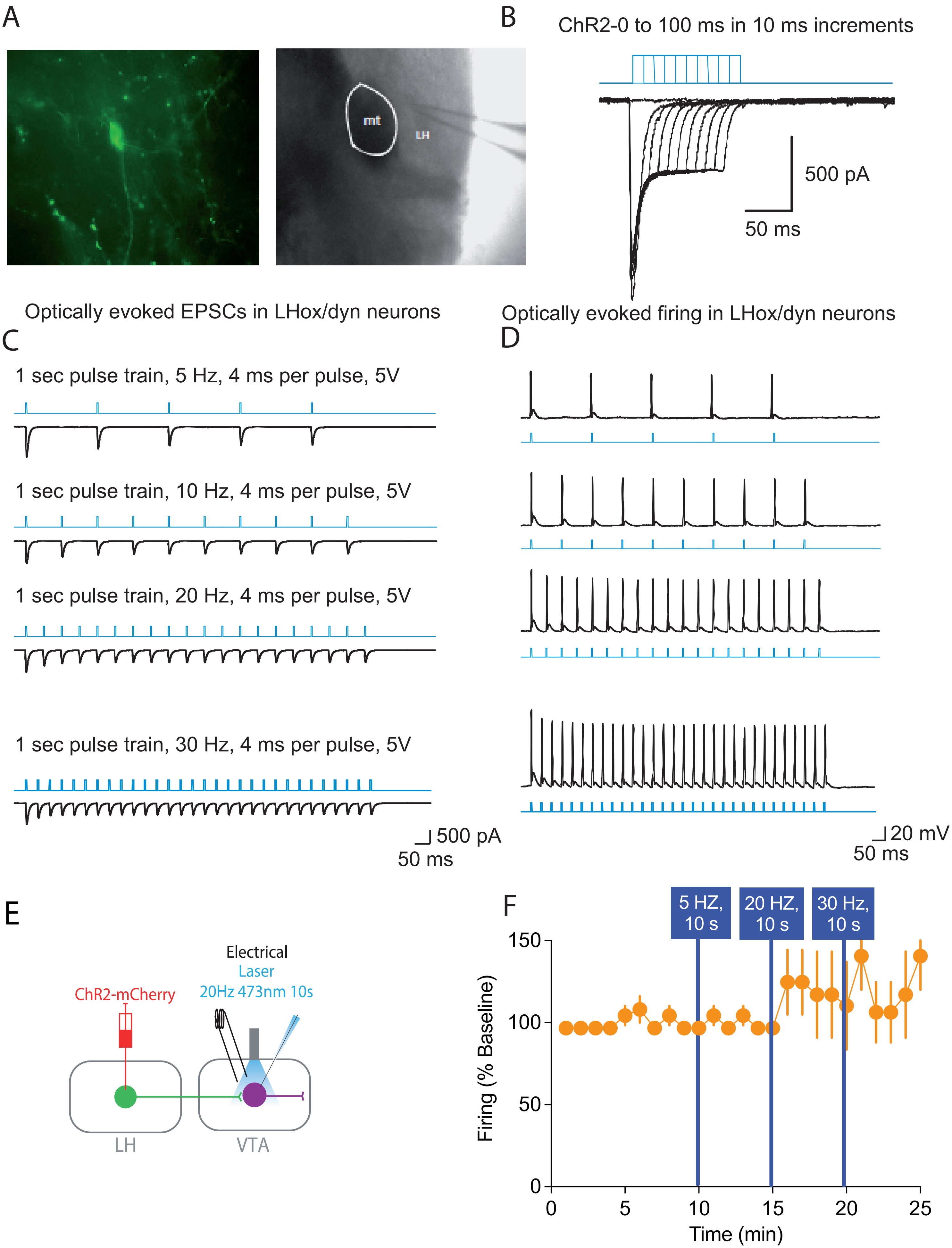


Supplemental Figure 1

### Supplemental Figure 2

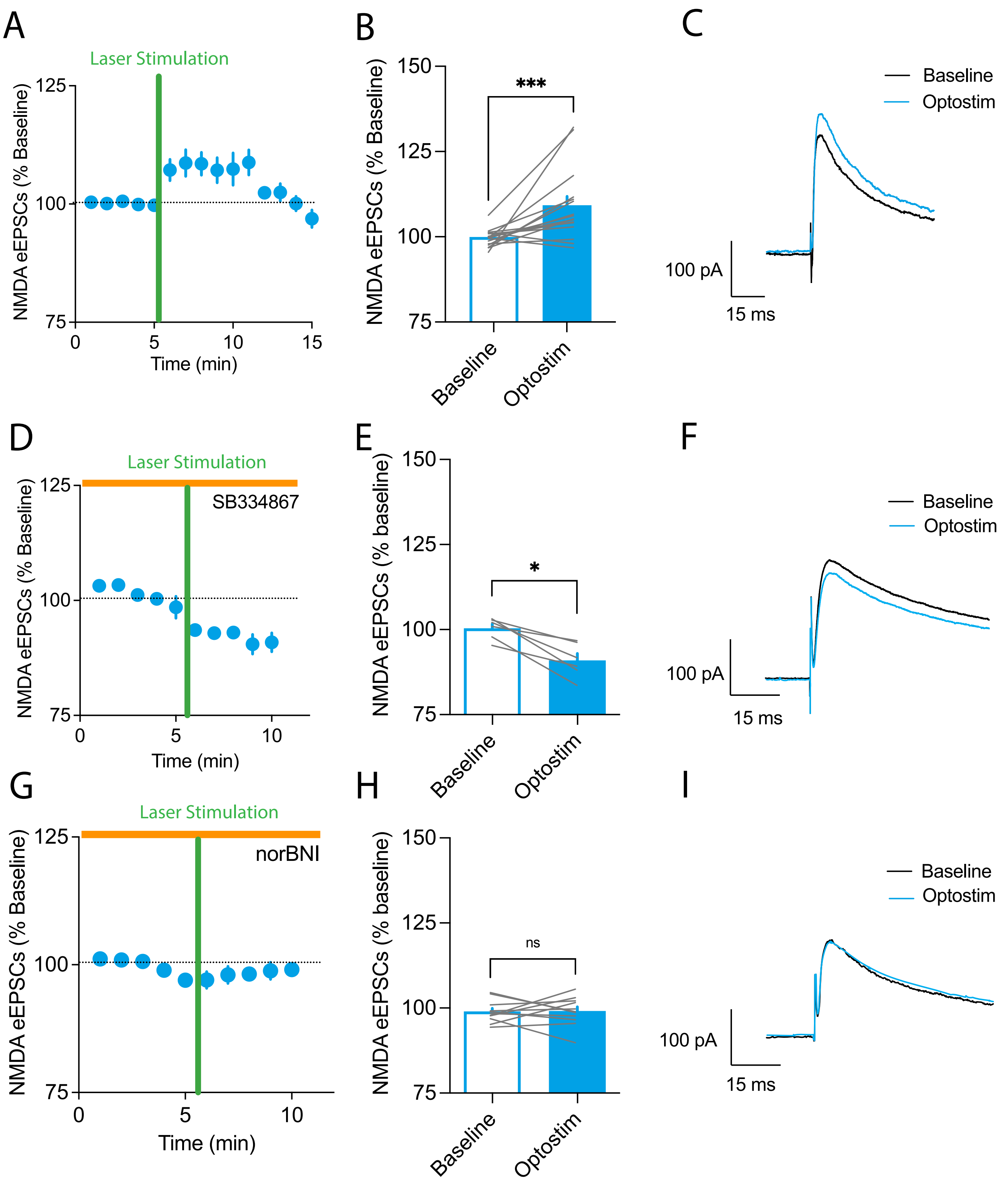
